## Supplementary Figures for "A New Gene Set Identifies Senescent Cells and Predicts Senescence-Associated Pathways Across Tissues"

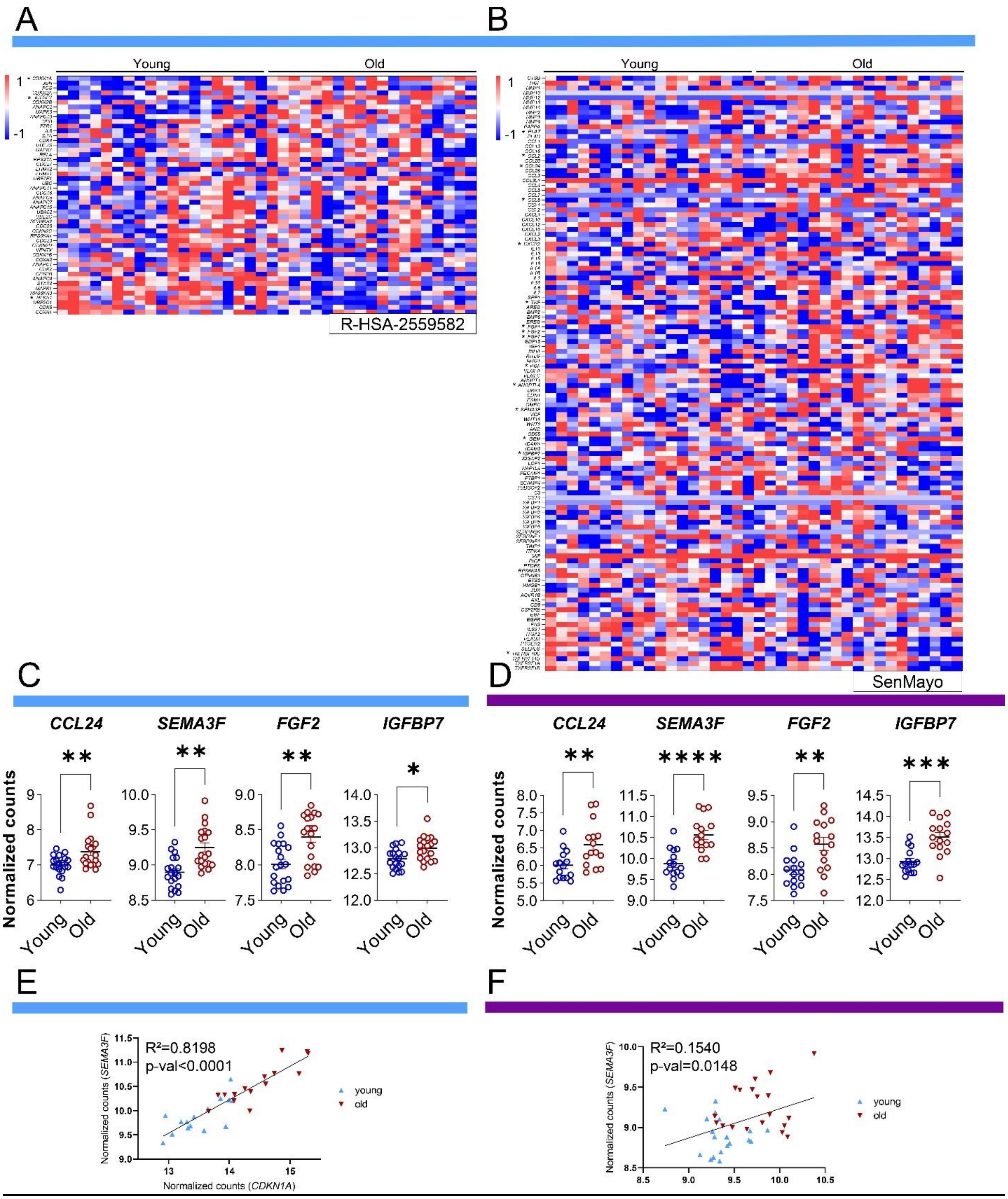


Figure S1. *The SenMayo gene set predicts aging in two mRNA-seq data sets.* Out of the 50 available genes in the R-HSA-2559582 gene set, two were significantly enriched in the aging cohort (A), while 13 out of 125 of the SenMayo genes were enriched (B). Canonical markers of the SASP such as *CCL24, SEMA3F, FGF2, and IGFBP7* were upregulated with aging in the RNAseq of human bone/bone marrow samples in cohort A (C) and cohort B (D). Moreover, the senescence markers, *CDKN1A/p21*$^{CIP1}$ and SEMA3F, correlate with each other in cohort A (E) and cohort B (F), demonstrating a potential circumvention of high interindividual variability by combining more than one gene. *p<0.05, **p<0.01, ***p<0.001, ****p<0.0001. Cohort A (blue): n=38 (19 young, 19 old, all ♀), Cohort B (purple): n=30 (15 young, 15 old, all ♀).

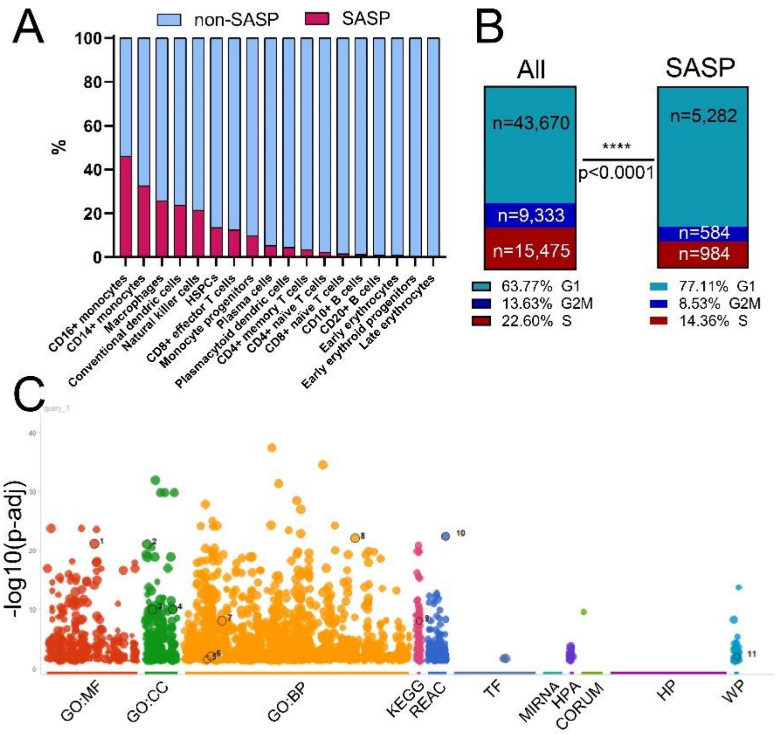


| id | source | term_id | term_name | term_size | | p_value |
|---|---|---|---|---|---|---|
| 1 | GO:MF | GO:0036094 | small molecule binding | 2506 | | 7.7e-22 |
| 2 | GO:CC | GO:0005615 | extracellular space | 3594 | | 9.6e-22 |
| 3 | GO:CC | GO:0030141 | secretory granule | 851 | | 1.1e-10 |
| 4 | GO:CC | GO:0099503 | secretory vesicle | 1020 | | 1.1e-10 |
| 5 | GO:BP | GO:0007050 | cell cycle arrest | 241 | | 3.2e-02 |
| 6 | GO:BP | GO:0008285 | negative regulation of cell population proliferation | 790 | | 8.8e-03 |
| 7 | GO:BP | GO:0010941 | regulation of cell death | 1720 | | 8.8e-09 |
| 8 | GO:BP | GO:0140352 | export from cell | 1418 | | 1.0e-22 |
| 9 | KEGG | KEGG:04911 | Insulin secretion | 86 | | 9.8e-09 |
| 10 | REAC | REAC:R-HSA-382551 | Transport of small molecules | 718 | | 3.9e-23 |
| 11 | WP | WP:WP254 | Apoptosis | 86 | | 9.6e-03 |

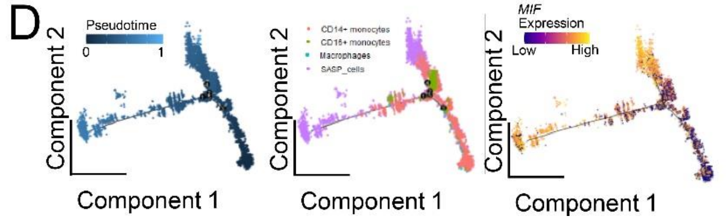


Figure S2. *The senescent phenotype and communication patterns of SASP cells in human hematopoietic bone marrow.* (A) 46% of $CD16^+$ monocytic cells had (high) SASP expression pattern; (B) The SASP cell cluster displayed a shift towards the G1 cell cycle phase, suggesting reduced replicative potential; (C) The enriched terms of the SASP cluster, depicted in a Manhattan plot, show the high expression of cell cycle arrest (GO: 0007050) and negative proliferation patterns (GO: 0008285); (D) SASP cells emerged in the final phases of cellular differentiation and increased *MIF* expression (yellow on the left, late phase) at their late developmental phase as revealed by pseudotime analysis. ****p<0.0001, n= 22 (10 ♂, 12 ♀).

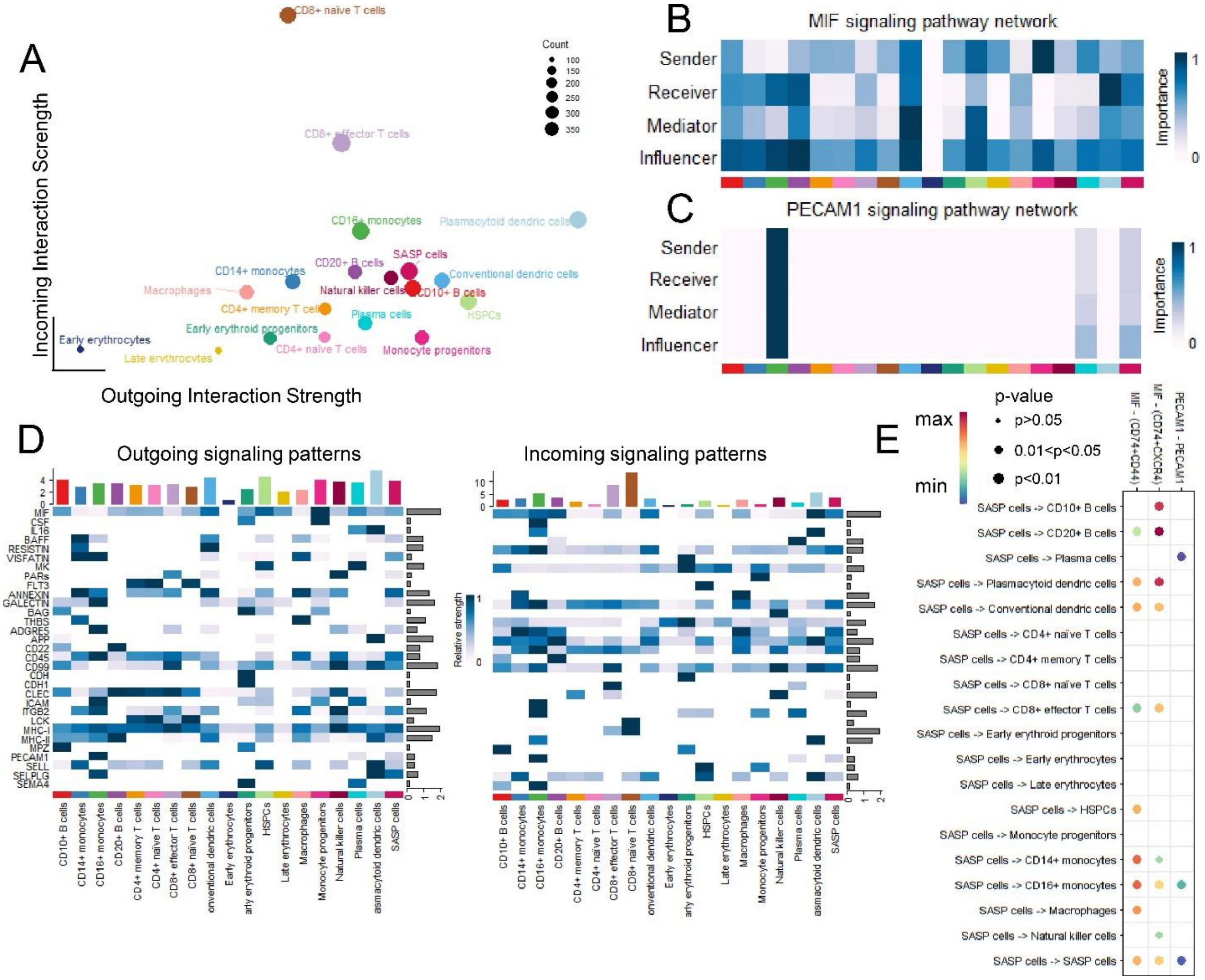


<u>Figure S3.</u> *Phenotypic origin of the SASP cluster in human hematopoietic bone marrow cells.* (A) The SASP cluster has an overall high outgoing and moderate incoming interaction strength; (B) The SASP cells exerted various signaling functions (sender, receiver, mediator, and influencer) in the MIF pathway and (C) the PECAM1 pathway, which was used mostly by CD16$^{+}$ monocytes, plasma cells, and SASP cells (color-code in D); (D) The outgoing signaling pattern revealed the relevance of the MIF pathway among all other pathways, while the relative incoming signaling pattern was likewise substantial; (E) A direct MIF-driven interaction (*via* CD74/CD44 or CD74/CXCR4) from the SASP cells was detected among the majority of other cell types, especially plasmacytoid dendritic cells and B cells, while the PECAM1 pathway mostly targeted the abovementioned three cell types, n= 22 (10 ♂, 12 ♀).

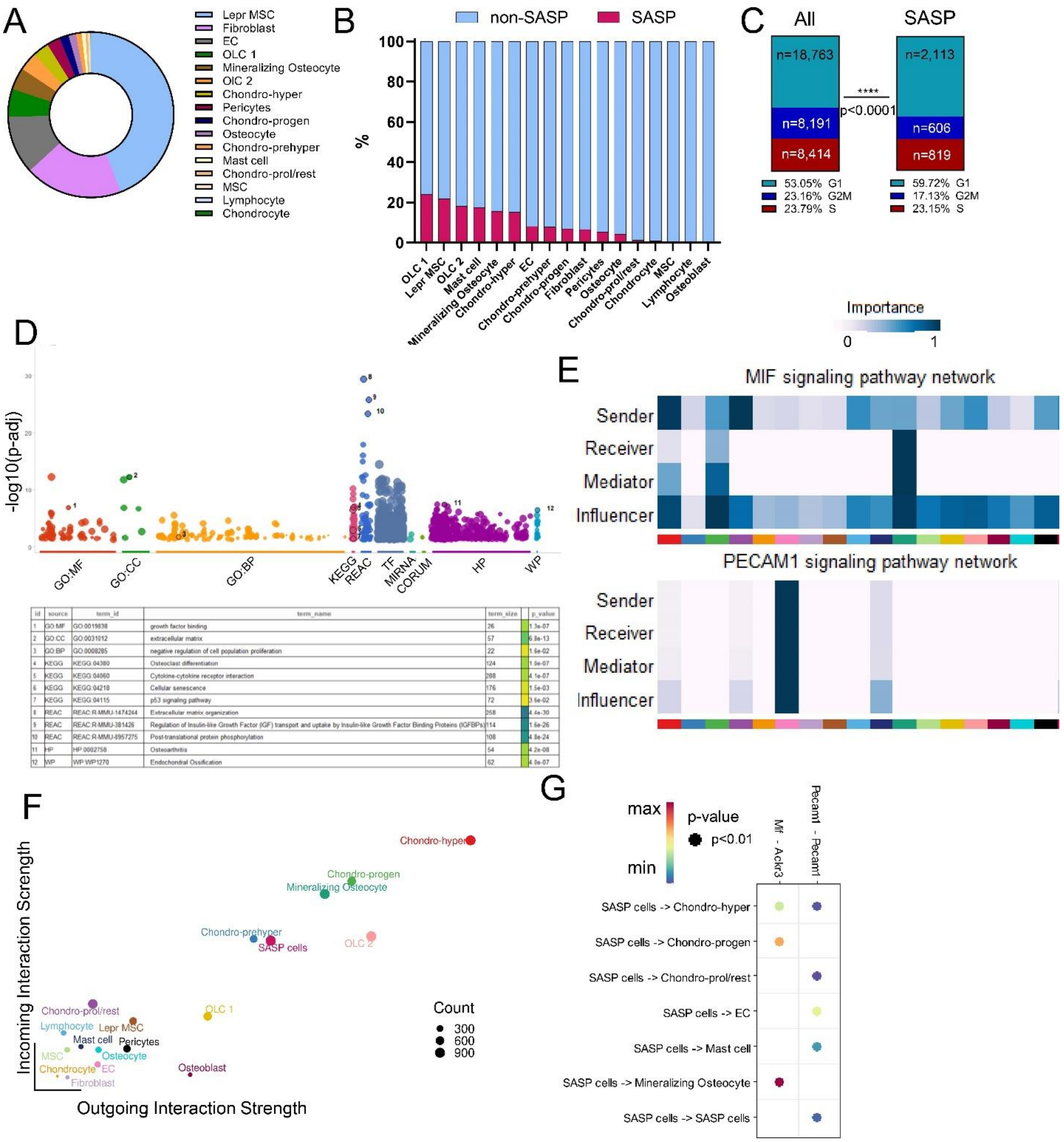


| id | source | term_id | term_name | term_size | p_value |
|---|---|---|---|---|---|
| 1 | GO:MF | GO:0019838 | growth factor binding | 26 | 1.3e-07 |
| 2 | GO:CC | GO:0031012 | extracellular matrix | 57 | 6.8e-13 |
| 3 | GO:BP | GO:0008285 | negative regulation of cell population proliferation | 22 | 1.6e-02 |
| 4 | KEGG | KEGG:04380 | Osteoclast differentiation | 124 | 1.0e-07 |
| 5 | KEGG | KEGG:04060 | Cytokine-cytokine receptor interaction | 288 | 4.1e-07 |
| 6 | KEGG | KEGG:04218 | Cellular senescence | 176 | 1.5e-03 |
| 7 | KEGG | KEGG:04115 | p53 signaling pathway | 72 | 3.6e-02 |
| 8 | REAC | REAC:R-MMU-1474244 | Extracellular matrix organization | 258 | 4.4e-30 |
| 9 | REAC | REAC:R-MMU-381426 | Regulation of Insulin-like Growth Factor (IGF) transport and uptake by Insulin-like Growth Factor Binding Proteins (IGFBPs) | 114 | 1.6e-26 |
| 10 | REAC | REAC:R-MMU-8957275 | Post-translational protein phosphorylation | 108 | 4.8e-24 |
| 11 | HP | HP:0002758 | Osteoarthritis | 54 | 4.2e-08 |
| 12 | WP | WP:WP1270 | Endochondral Ossification | 62 | 4.0e-07 |

Figure S4. *Murine SASP cells in mesenchymal cells from bone and bone marrow are mainly of osteolineage origin.* (A) SASP cells were mostly recruited from osteolineage cells (OLC), and leptin-positive ($Lep^+$) mesenchymal stem cells (MSCs), while (B) 24% of OLC 1 and 18% of OLC2 cells were SASP cell members; (C) Mesenchymal SASP cells in murine bone and bone marrow significantly changed their replicative state from G2M to G1, indicating a replicative stop; (D) A Manhattan plot depicts an enrichment of genes associated with cellular senescence (KEGG 04218), negative regulation of proliferation (GO0008285), and cytokine-receptor interaction (KEGG 04060) within the SASP cluster; (E) SASP cells function as both senders and influencers within the *MIF* network, and mostly as influencers in the PECAM1 network; (F) The outgoing interaction strength of the SASP cells was high, while they simultaneously showed a substantial incoming signaling strength; (G) Direct cell-cell interactions in the *MIF* pathway from the SASP cells is predominantly directed to hypertrophic chondrocytes, chondrocytic progenitors, and mineralizing osteocytes, while the Pecam1 pathway is directed to chondrocytes, endothelial cells, mast cells, and the SASP cells themselves. ****$p<0.0001$, n= 8 (4 bone, 4 bone marrow, all ♂).

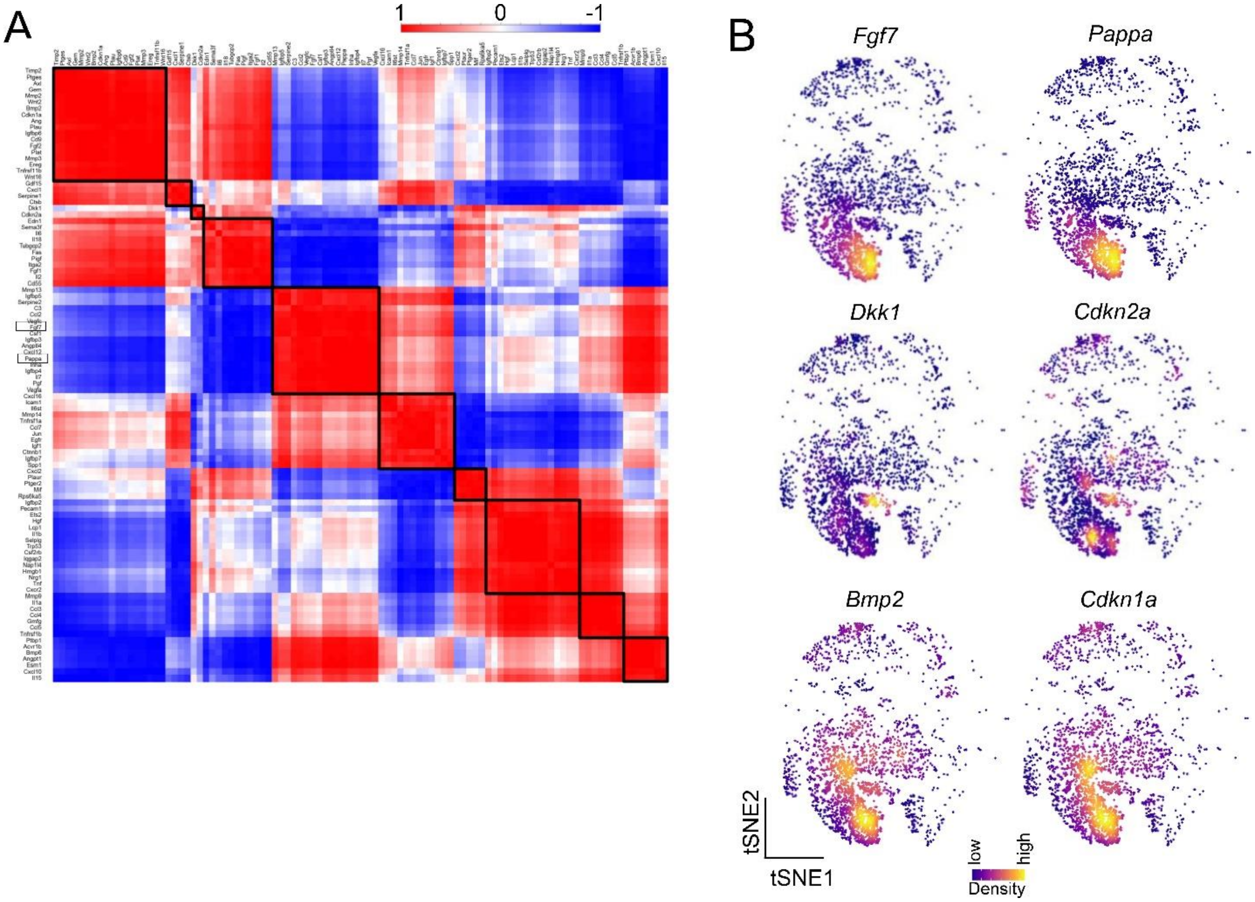


Figure S5. *The murine SASP cluster showed an enrichment of senescent pathways and contained distinct expression modules.* (A) Within the SASP cluster, different expression modules were mathematically predicted. In this module, *Pappa* and *Fgf7* are present, which can be visualized spatially in (B) tSNE, having a similar kernel-weighed density. Other predicted co-expressional patterns were demonstrated by the pairs *Dkk1-Cdkn2a* and *Bmp2-Cdkn1a*. n= 8 (4 bone, 4 bone marrow, all ♂).
